# Shared responsibility decreases the sense of agency in the human brain

**DOI:** 10.1101/2021.08.20.457115

**Authors:** Marwa El Zein, Ray J. Dolan, Bahador Bahrami

**Affiliations:** Center for Adaptive Rationality, Max Planck Center for Human Development; Berlin, Germany; Institute of Cognitive Neuroscience, University College London; London, United Kingdom; Max Planck University College London Centre for Computational Psychiatry and Ageing Research, University College London; London, United Kingdom; Wellcome Centre for Human Neuroimaging, University College London; London, United Kingdom; Faculty of Psychology and Educational Sciences, Ludwig Maximilian University; Munich, Germany; Department of Psychology, Royal Holloway, University of London; Egham, Surrey, United Kingdom

## Abstract

Sharing responsibility in social decision-making helps individuals use the flexibility of the collective context to benefit themselves by claiming credit for good outcomes or avoiding the blame for bad outcomes. Using magnetoencephalography (MEG), we examined the neuronal basis of the impact that social context has on this flexible sense of responsibility. Participants performed a gambling task in various social contexts and reported feeling less responsibility when playing as a member of a team. A reduced MEG outcome processing effect was observed as a function of decreasing responsibility at 200 ms post outcome onset and was centred over parietal and pre-central brain regions. Prior to outcome revelation in socially made decisions, an attenuated motor preparation signature at 500 ms after stimulus onset was found. A boost in reported responsibility for positive outcomes in social contexts was associated with increased activity in regions related to social and reward processing. Together, these results show that sharing responsibility with others reduces agency, influencing pre-outcome motor preparation and post-outcome processing, and provides opportunities to flexibly claim credit for positive outcomes.

## Introduction

In collective decision-making, we have less control over the choices and outcomes than when we are making decisions alone. We are, however, not completely bound by constraints or instructions as to what to do. In return for this partial concession of control to the collective, we benefit from a *sharing* of responsibility for our choices [1]. Indeed, when people assign credit to contributors in a team, they tend to overestimate their own contribution and over attribute a team’s success to their own abilities, effort and merit [2]. Conversely, when outcomes are poor, the collective context allows us to distance ourselves from regret [3] and offload blame onto others [4]. Teams are more likely to violate rules than individuals [5] and, correspondingly, people find it harder to punish groups (vs individuals) that have violated a social norm [4]. The advantages of such flexibility are not restricted to the psychology laboratory and can be observed in everyday life. When weapons of mass destruction were not found in Iraq in 2003 or the years that followed, intelligence agencies whose reports had justified the catastrophic invasion of Iraq defended themselves by claiming that “*everyone had agreed at the time*”.

The brain mechanisms that underly our sense of responsibility for the outcomes of our actions have generally been investigated by comparing active vs forced (or passive) choices [6–10]. The subjective experience of a coerced (or instructed) action is similar to that of a passive action and is associated with reduced neural processing of an action’s outcome [6]. These studies have invariably focused on the context of isolated individuals making private decisions. However, in social contexts, e.g., voting, we do not operate in the extremes of free vs coerced choice. As explained above, the collective context affords a level of cognitive flexibility that helps individuals favourably serve themselves by claiming credit or avoiding blame. As such, examining the neurobiological basis of shared responsibility and agency in the collective context opens a unique and novel door to the flexibility of human cognition that goes beyond earlier studies on the neurobiology of agency in private decision-making.

In this study, we investigated the neurobiological substrates of this flexible sense of responsibility in a social collective context. Operationally, we defined responsibility as a participant’s subjective judgement on the causal attribution between a decision and its outcome. In this sense, we followed the lead of earlier literature that proposes a strong connection between responsibility, the ‘sense of agency ’and feeling of control [1,6,11,12]. We replicated previous investigations that focused on the evaluation of outcomes in free vs instructed decision-making by individuals [6,13–15] and went beyond those studies to examine the impact of various collective group sizes on the processing of positive and negative outcomes.

Moreover, it has been suggested that examining the brain’s responses to outcomes has provided a very convenient methodological approach to the complicated concept of responsibility [12]. Our experimental design permitted us to go one important conceptual step further and ask if the neurobiological substrates of responsibility in the human brain emerge *during* deliberation, thus *before* a choice is made and the outcome is known.

In a choice-based gambling task, participants made decisions that led to positive or negative outcomes, while responsibility was parametrically modulated by the impact of different social contexts. We constructed four different contexts: (1) *Private*, in which the individual participant assumed full responsibility; (2) *Dyadic* and (3) *Group*, in which the individual decided together with one or four other people, respectively, and shared the responsibility with them; and (4) *Forced*, where another person decided on behalf of the participant, thus absolving the participant of all responsibility. Critically, the statistical frequencies of various outcomes were kept constant across all conditions, thus controlling for the expected value of options and choices. Another important issue that distinguishes our design from previous works is the distinction between actions and decisions. A number of previous studies showed that performing *actions* together with others reduces subjective ratings of responsibility and control [14,16,17]. However, our study is the first to investigate joint responsibility for collective *decisions*. We expected responsibility to be highest in the *Private* context and progressively decrease from *Dyadic* to *Group*, and then to its lowest level in the *Forced* context (behavioural **Hypothesis 1a**).

We used magnetoencephalography (MEG), which provides a high temporal resolution neural signal, to unravel the dynamics in the neural processes that underly responsibility in social contexts at various stages of the task. Our design allowed us to conduct trial-by-trial regressions of responsibility contexts with MEG signals [18,19], instead of comparing grand-average, event-related fields under high vs low responsibility contexts as previously done [6,13–15]. Previous studies found a decrease in the neural signatures of outcome processing that resulted in reduced responsibility, for example, when people are coerced to perform an action (vs willingly performing the same action)—an auditory tone signalling the outcome of the action evoked a lower, auditory, evoked potential (N1) [6]. Moreover, the feedback related negativity (FRN)—an evoked potential that appears around 200 to 300 ms after outcome onset—is also attenuated when an outcome resulted from an action performed in the presence of another agent [13], with others [14], and during a task where participants’ control over outcomes was modulated [15]. A fronto-parietal brain network—including the inferior parietal lobule, the angular gyrus, and the premotor and motor cortices—is implicated in the subjective feeling of control [7,20–22]. Based on these findings, we predicted attenuated outcome processing as a function of decreased responsibility, in particular at 200-300 ms after outcome onset with localisation in a fronto-parietal network (**Hypothesis 2**).

To identify neural signatures of outcome-independent, prospective responsibility, we examined a MEG signal for a sense of agency in the time period prior to the outcome. It has been suggested that when deliberating on an action, our sense of agency stems from a mental simulation of that action [12,23,24]. Prospective responsibility, in this sense, corresponds to the mental simulation of the likely outcomes of imagined actions. One study, in which participants underwent functional Magnetic Resonance Imagery (fMRI) while reading vignette scenarios and imagining themselves or others as the protagonist, showed that the contemplation of the consequences of imagined actions recruited premotor cortex [25] consistent with a mental simulation account. Thus, we predicted that prior to an outcome, the preparation of motor activity would be modulated by the responsibility context. Specifically, we hypothesised that lateralized motor preparation signals [18,26] would be attenuated in social contexts where responsibility is reduced (**Hypothesis 3**).

Finally, we addressed the influence of outcome valence and its interaction with responsibility. Previous research has shown that people tend to have a self-serving bias, whereby they take more credit for positive outcomes as compared to negative outcomes— an effect evident in both group and individual decision-making [27–31]. Based on this data, we expected to observe increased responsibility ratings for positive vs negative outcomes irrespective of whether participants made decisions alone or with others, but not when they had no responsibility for the decision, i.e., in the Forced context (**Hypothesis 1b**). In addition, we hypothesised that, since social contexts offer a possibility to share responsibility, this would enable participants to cherry pick credit for positive outcomes and offload blame for negative outcomes onto others [1], thereby exaggerating a self-serving bias in the Dyadic and Group contexts vs the Private context. Importantly, our experimental design allowed us to examine two distinct possible mechanisms: a concurrent increase of credit and decrease in blame, or a specific modulation of either credit or blame respectively (**Hypothesis 1c**).

## Results

### Behavioural results

Participants performed a task where they had to choose between two gambles that could yield positive or negative outcomes. They did so in four different contexts: Alone (*Private context*), with one other player (*Dyadic context*), in a group of five players (*Group context*), and where another player decided for them (*Forced context*) (**Figure 1a**). In the last context, Forced, participants were requested to press the right or left button even though it did not count as their choice. As they already knew which button to press, participants were faster to respond in a Forced context compared to all the other contexts (all T_39_>6.4, all p<0.001). Participants were also faster at making a decision in the Dyadic context as compared to the Group context (T_39_=-2.63, p=0.01, μ=-15.94, ci=[−28.18, −3.71], dz=0.41), but no other differences were observed. After a decision was made between the two gambles, participants used a scale to report how responsible they felt for the outcome of the trial. These responsibility ratings represent our main behavioural variable of interest (**Figure 1b**).

**Figure 1.**
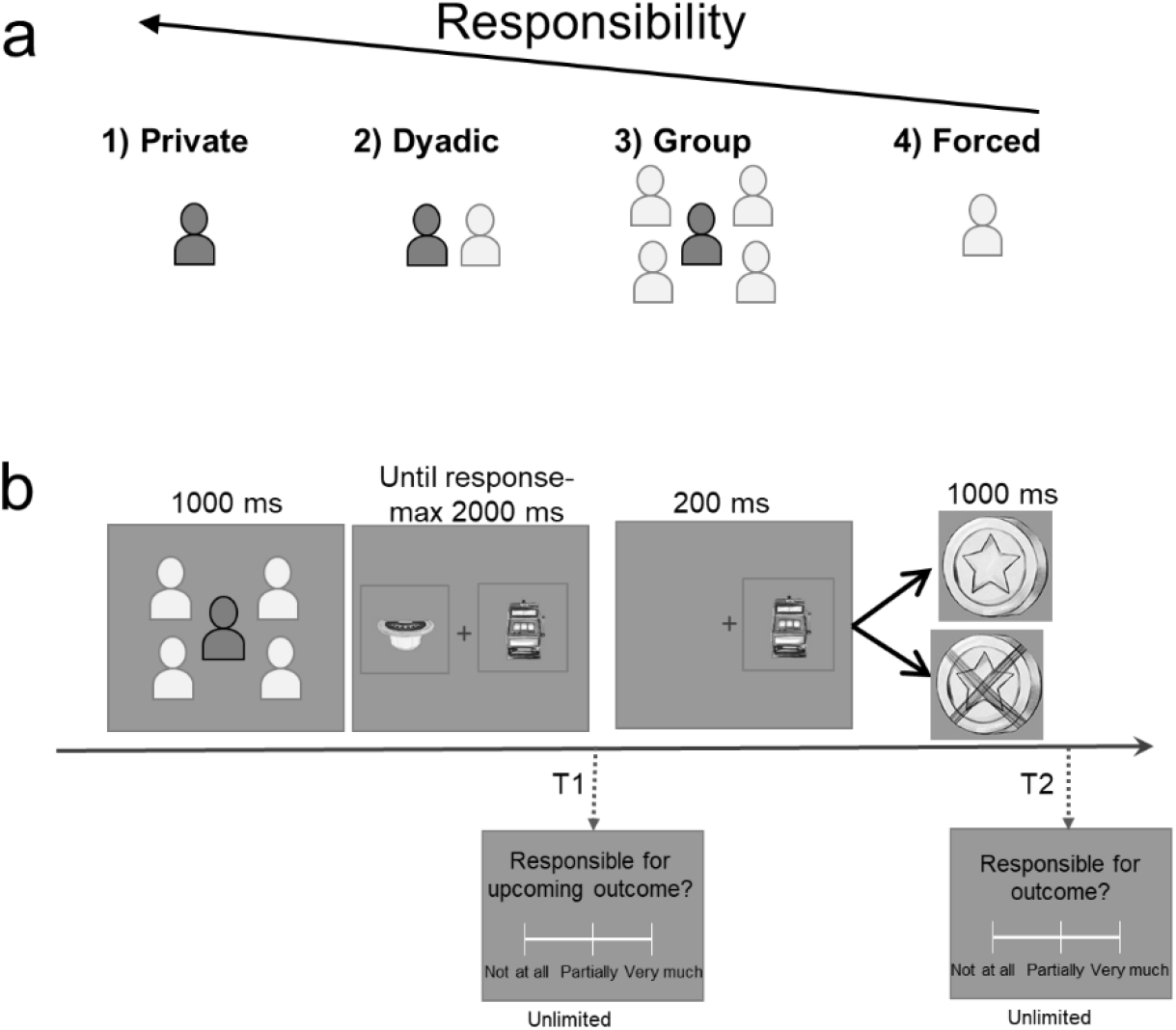
Experimental design. a) Participants performed a decision-making task in four contexts: (1) Private: playing alone; (2) Dyadic: together with another participant; (3) Group: together in a group of five participants; and (4) Forced: another participant played for them. Dyadic and Group contexts are henceforth referred to as Social contexts. b) Each trial (384 trials in 8 blocks), began with participants having 1 second to see the context (as described in *a*) in which they would be playing. Next, they chose between two gambling options that were displayed on the screen for a maximum of 2 seconds. After making a choice, the selected gamble remained on the screen for 200 ms. In the Social contexts, the selected gamble may or may not coincide with the gamble chosen by the group. Finally, the outcome (positive or negative) was displayed for 1 second. The trials of each block were randomly assigned into three groups. One-third of participants rated how responsible they felt for the upcoming outcome on a continuous scale immediately after the gamble selection (here marked by T1). In another third of the trials, participants rated how responsible they felt for the obtained outcome on a continuous scale after the outcome (T2). In the remaining third of the trials, no rating was required.

#### Parametric responsibility reporting

To test **Hypothesis 1a**, we regressed z-scored, continuous responsibility ratings against responsibility contexts (1= Private, 2= Dyadic, 3= Group, 4= Forced) for each participant. All participants except for one (p>0.2) showed a significant negative slope (37 participants p<0.001, 1 participant p<0.005, 1 participant p<0.02), i.e., they reported a linearly decreasing sense of responsibility, starting from the Private context and moving down to the Dyadic, Group and then Forced contexts (t-test of the betas computed for each participant against zero, T_39_=-19.72, p<0.001, μ =-0.61, ci=[−0.67,−0.55], dz=-3.11) (**Figure 2a**). T-tests comparisons between contexts confirmed this linear change in responsibility: Responsibility ratings were higher in the Private context as compared to the Dyadic context (T_39_=12.14, p<0.001, μ =0.76, ci=[0.63,0.89], dz=1.95). Dyadic context ratings were higher than Group context ratings (T_39_=6.08, p<0.001, μ =0.24, ci=[0.16,0.33], dz=0.96). Finally, Group context ratings were higher than those in the Forced context (T_39_=13.19, p<0.001, μ =0.96, ci=[0.81,1.10], dz=2.08).

**Figure 2.**
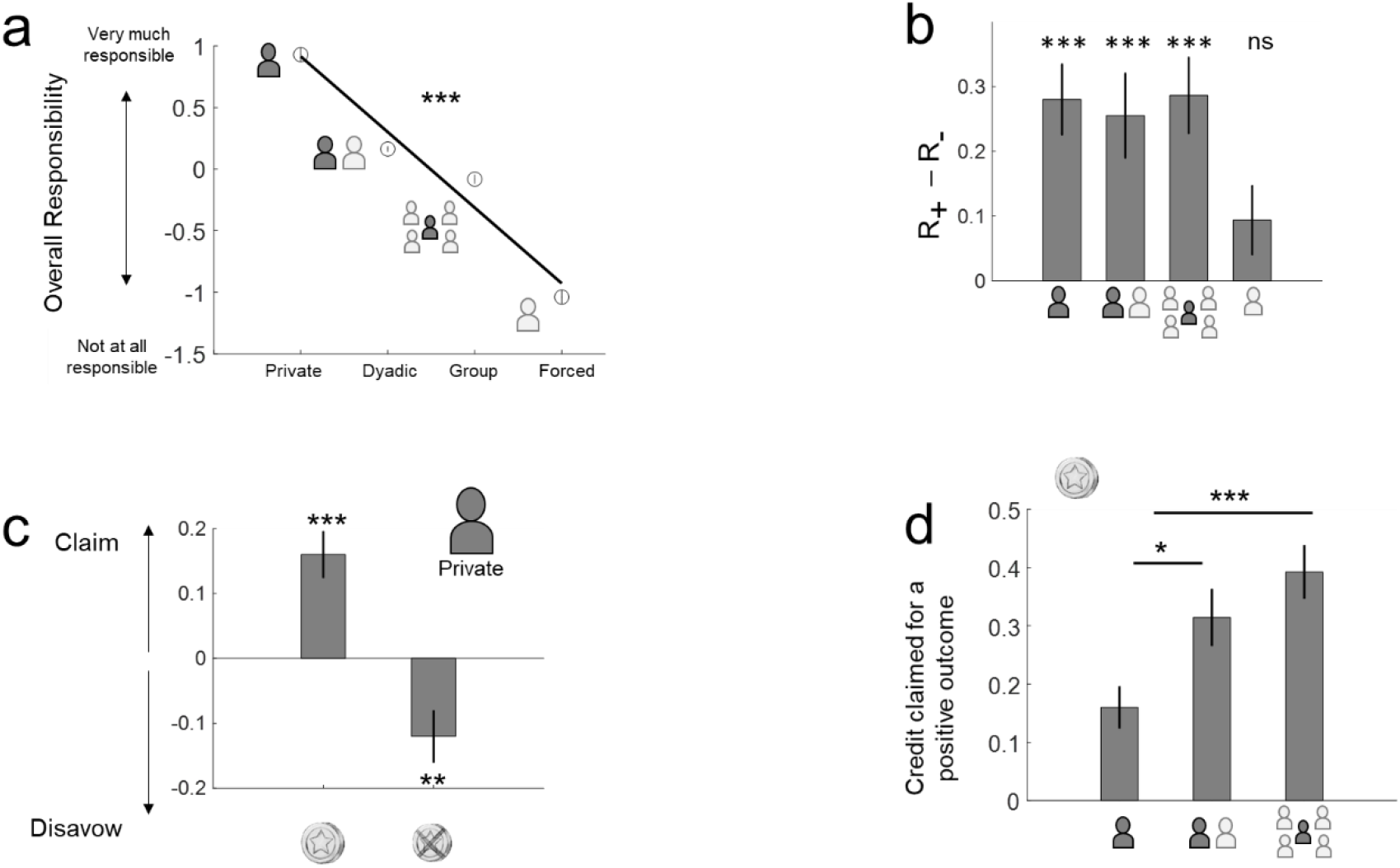
Behavioural results. Responsibility ratings were reported on a continuous scale from ‘not at all responsible ’to ‘very much responsible ’(see **Fig. 1b**). The ratings were z-scored for each participant for the analyses and are therefore reported in arbitrary units in all graphs. (a) Parametric decrease of reported responsibility from Private to Dyadic to Group to Forced contexts. Circles represent the mean values across participants for each context with the associated standard errors. The line is the mean general linear model fit (fit for each participant, then averaged across all 40 participants). (b) The impact of outcome valence on reported responsibility. The Y-axis shows the differences in responsibility claimed for positive (R_+_) and negative (R_−_) outcomes. In all cases where the participant had *some* choice in the selection of the gamble, they claimed more responsibility for positive (vs negative) outcomes. The Forced context, where the subject had no say in gamble selection did not show a similar bias. (c) When compared to reported responsibility *before* the outcome, we observed that outcome valence modulated responsibility in both positive and negative directions. Data from the Private context alone: Participants both claimed more credit after positive outcomes and disavowed a negative outcome. The Y-axis shows the difference between responsibility ratings after vs before the outcome. The X-axis shows outcome valence. (d) When compared with pre-outcome ratings, people claimed more credit for a positive outcome in the social contexts when their vote was selected compared to the Private context. Error bars represent standard errors. ***: *p*<0.001, **: *p*<0.01, \**p*<0.05, *ns*: non significant.

These behavioural results confirm a reduced sense of responsibility in Social contexts, and furthermore show that this reduction depends on group size. Moreover, the reports of decreased responsibility validate our experimental paradigm, which was designed to address the neural processes involved in decision-making in different responsibility contexts.

#### Self-serving bias

Our behavioural **Hypothesis 1b** stated that responsibility ratings would reveal a self-serving bias, with participants attributing more responsibility to themselves for positive (vs negative) outcomes. We observed that participants indeed provided higher responsibility ratings for positive (vs negative) outcomes in the three active contexts (Private context: T_39_=5.18, p<0.001, μ =0.28, ci=[0.17,0.39], dz=0.82; Dyadic context: T_39_=3.92, p<0.001, μ =0.25, ci=[0.12,0.39], dz=0.61; and Group context: T_39_=4.89, p<0.001, μ =0.29, ci=[0.16, 0.40], dz=0.77), but not in the Forced context (T_39_=1.76, p>0.08, μ =0.09, ci=[-0.01,0.20], dz=0.27) (**Figure 2b**). The magnitude of this bias did not differ across the three active contexts (all p>0.48, all T>0.69). Contrary to our **Hypothesis 1c**, we found no evidence for an increase in self-serving bias in the Social (vs Private) contexts.

#### Claiming credit for success or disavowing blame for failure?

Next, we examined if a self-serving bias was observed through the attribution of credit after positive outcomes or disavowal of responsibility after negative outcomes. In different trials, we asked participants to report their responsibility ratings *before* and *after* an outcome. Taking the *before* ratings as a baseline, we then assessed whether the self-serving bias consisted of higher ratings after positive outcomes and/or lower ratings after negative outcomes. For each outcome valence, we subtracted the *before* ratings from the *after* ratings.

In the Private context, a self-serving bias was demonstrated in both components: more responsibility was claimed after (vs before) positive outcomes (T_39_=4.41, p<0.001, μ=0.16, ci=[0.09,0.23], dz=0.72) and less responsibility was accepted after (vs before) negative outcomes (T_39_=-2.96, p<0.01, μ =-0.12, ci=[-0.20,0.03], dz=-0.46) (**Figure 2c**).

In the Social contexts, a more complex analysis was required to accommodate the relationship between a participant’s decision, the collective choice and the outcome. When the collective choice matched the participant’s decision, responsibility ratings were boosted after (vs before) an outcome (T_39_>4.26, p<0.001). It is important to note that participants claimed more credit in Social contexts (vs the Private context) for positive outcomes (Private vs Dyadic: T_39_=-2.51, p<0.02, μ =-0.15, ci=[-0.27,-0.03], dz=-0.39; Private vs Group: T_39_=-4.39, p<0.001, μ =-0.23, ci=[-0.33,-0.12], dz=-0.70) (**Figure 2d**). This finding is partly consistent with **Hypothesis 1c** in revealing how Social contexts offered a ‘cover ’for claiming more credit for a positive outcome. Participants, however, did not disavow responsibility for negative outcomes (T_39_>0.59, p<0.55), which led to a conclusion that was opposite to our prediction: participants disavowed negative outcomes more in Private vs Social contexts (Private vs Dyadic: T_39_=-3.05, p=0.004, μ =-0.15, ci=[-0.25,-0.05], dz=-0.48; Private vs Group: T_39_=-2.79, p=0.007, μ =-0.14, ci=[−0.25,−0.04], dz=−0.42). In trials where the collective choice was different from the participant’s decision, responsibility ratings were generally lower after (vs before) an outcome (T_39_>5.3, p<0.001) (see *Supplementary material* for a full description of these results).

### MEG results

#### Outcome processing is parametrically modulated by shared responsibility

Our key neural hypothesis stated that neural signatures for shared responsibility would be common to those identified for a sense of agency. **Hypothesis 2** stated that outcome processing within 200-300 ms after outcome onset would vary linearly with responsibility levels. This signal would locate to a fronto-parietal brain network that is associated with a sense of agency and includes the inferior parietal lobule, angular gyrus, and the premotor and motor cortices. We isolated pre-processed MEG signals in this time window and performed whole-brain regressions of those signals against the responsibility contexts (see *Methods*). Two clusters of electrodes showed significant activity which survived correction for multiple comparisons (cluster alpha < 0.05; **Figure 3a,** right panel): these electrodes match an expected fronto-parietal network, in that they predominantly consist of central, frontal and parietal electrodes (see full list of electrodes in *Supplementary materials*). The mean parameter estimate of the regression for the significant right-side electrodes is shown through time in Figure 3a, left panel (statistics of the effect at its peak in the 200-300 ms window at 220 ms (T_39_=-4.61, p<0.001, μ =−4.15, ci=[−5.98,−2.33], d=0.72). With these same right-side electrodes, we performed an additional descriptive analysis, computing event-related fields locked to outcome, separately for each context. The largest event-related field was observed for the Private context and then decreased linearly through the Forced context at ~200 ms after outcome onset (**Figure 3b**). This is in line with the trial-by-trial GLM results. Finally, the same GLM regressions in the source space (see *Methods*) revealed that the parametric encoding of responsibility at 200-300 ms after outcome onset is associated with parietal and central sources (**Figure 3c**), which is described in detail in Table 1.

**Figure 3.**
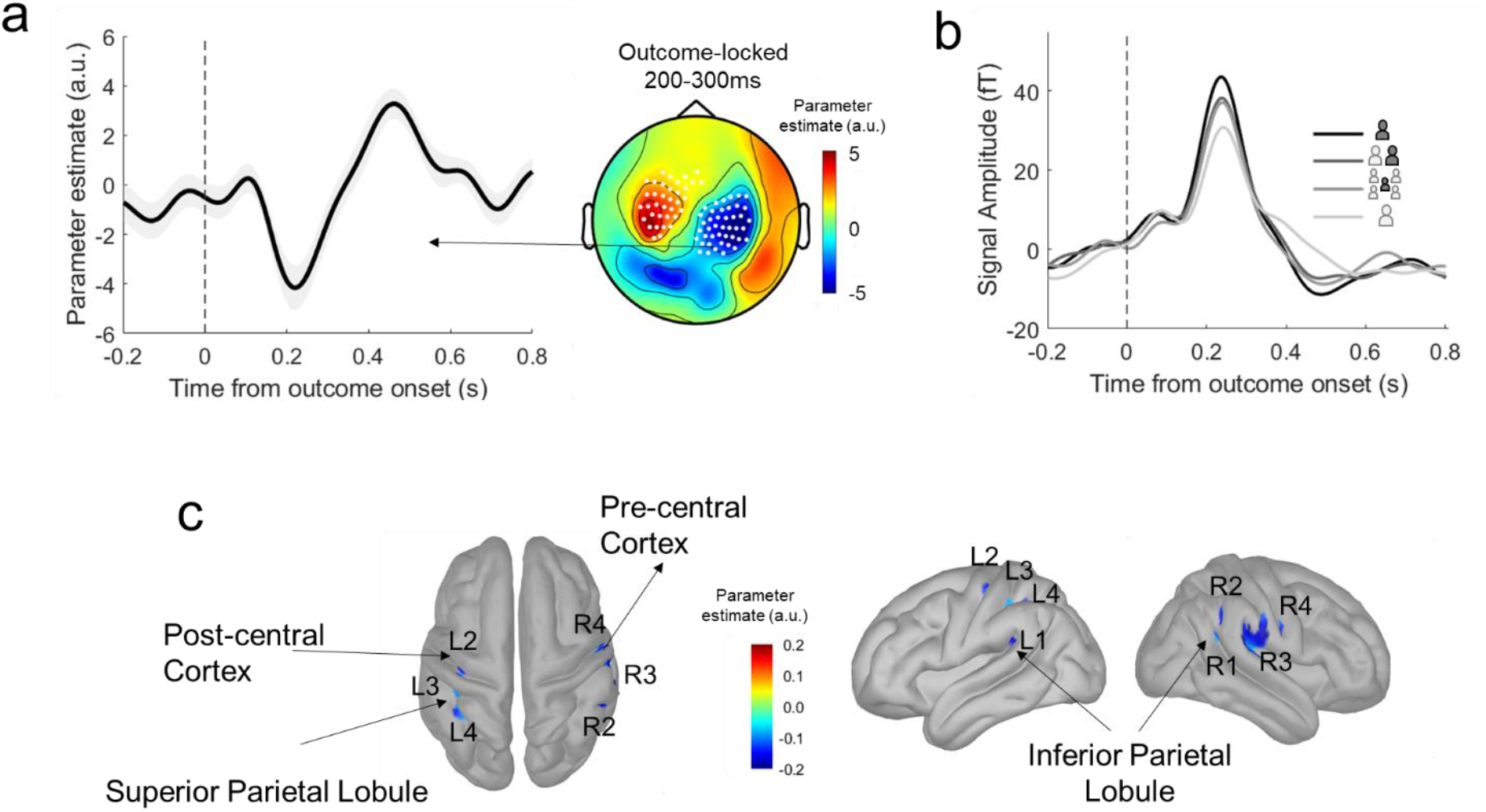
Agency-related neural correlates of responsibility at 200 to 300 ms after outcome onset. a) Parameter estimate of responsibility regression. Right panel: Scalp topography of the parameter estimate of responsibility regression [1 Private, 2 Dyadic, 3 Group of Five, 4 Forced] on the mean MEG activity at 200-300 ms after outcome. White dots represent the significant electrodes where the MEG signal linearly co-varies with the level of responsibility using cluster corrections of the effect against zero at an alpha cluster level <0.05. Left panel: Parameter estimate of responsibility regression at the right cluster locked to outcome onset. b) Associated Event-Related Fields (ERF) at the same right electrode cluster show how the amplitude of the ERF locked to outcome increases with responsibility. c) Estimated cortical sources of the responsibility parameter estimate at 200-300 ms. Parameter estimates of the responsibility regression at the source level that are significant at p<0.01 are shown. R=Right hemisphere and L=Left hemisphere, and the associated numbers refer to the different brain regions reported and anatomically localised in Table 1.

**Table 1.** Anatomical sources of parametric responsibility encoding locked to outcome. Regions were determined based on the Destrieux Atlas and implemented in Brainstorm. Then, MNI coordinates were extracted from Brainstorm and projected onto the BNA atlas.

|  | Destrieux Atlas |  |  | MNI coordinates |  |  | Human Brainnetome Atlas (BNA) |  |  |  |
| --- | --- | --- | --- | --- | --- | --- | --- | --- | --- | --- |
| Regions | Index | Name | Localisation | X | Y | Z | Label ID | Gyrus | Anatomy | Lobe |
| Left 1 (L1) | 41 | Posterior segment of the lateral sulcus | Insula | -48 | -40 | 25 | 145 | Inferior parietal lobule | Rostroventral area 40 | Parietal |
| Left 2 (L2) | 29 | Precentral gyrus | Lateral frontal lobe | -41 | -24 | 57 | 155 | Post central Gyrus | Area 1/2/3 |  |
| Left 3 (L3) | 67 | Postcentral Sulcus | Lateral Parietal Lobe | -34 | -36 | 42 | 129 | Superior Parietal Lobule | Lateral area 5 |  |
| Left 4 (L4) | 56 | Intraparietal sulcus and transverse parietal sulci |  | -34 | -45 | 39 | 129 |  |  |  |
| Right 1 (R1) | 73 | Superior temporal sulcus | Temporal lobe | 46 | -44 | 24 | 144 | Inferior parietal lobule | Rostroventral area 39 |  |
| Right 2 (R2) | 55 | Sulcus intermedius primus (of Jensen) | Inferior parietal lobule | 53 | -43 | 35 | 142 |  | Caudal area 40 |  |
| Right 3 (R3) | 26 | Supramarginal Gyrus |  | 58 | -19 | 17 | 146 |  | Rostroventral area 40 |  |
|  | 67 | Post Central Sulcus | Main sulci of the lateral aspect of the parietal lobe | 54 | -15 | 35 | 160 | Post central Gyrus | Area 2 |  |
|  | 36 | Temporal plane of the superior temporal gyrus | Superior aspect of the temporal lobe | 65 | -26 | 16 | 76 | Superior temporal Gyrus | Caudal area 22 | Temporal |
| Right 4 (R4) | 45 | Central Sulcus | Frontal Lobe | 51 | -5 | 25 | 54 | Precentral Gyrus | Area 4 | Frontal |

#### Neural correlates of a *prospective* (outcome-independent) sense of responsibility

Having established a neural signature of responsibility in outcome processing, we then investigated a neuronal expression of pre-outcome responsibility under Private and Social contexts. Here, we refer to responsibility experienced prior to choice and outcome, which we hypothesised would be related to motor preparation activity and subject to modulation by the responsibility contexts. Specifically, we tested whether lateralized motor preparation signals to select a gamble with the left or right hand [26] decreased in the Social contexts where responsibility is reduced. We first computed motor preparation signals at 100 ms before the response button was pressed by subtracting the power in the 8-32 Hz frequency band when participants responded with the right hand from when they responded with the left hand, which allowed us to identify the central electrodes with maximal suppression (**Figure 4a**, top panel). Then, we subtracted the power in the 8-32 Hz frequency band in ipsilateral minus contralateral maximal central electrodes relative to the hand pressed (**Figure 4a**, bottom panel). We found that motor preparation signals increase gradually until response in all four contexts. It should be noted that motor preparation signals in the Forced context, where the choice of button press was already known and the reaction times were fastest, diverged radically from the other conditions and were excluded from further hypothesis testing (see *Supplementary materials*).

**Figure 4.**
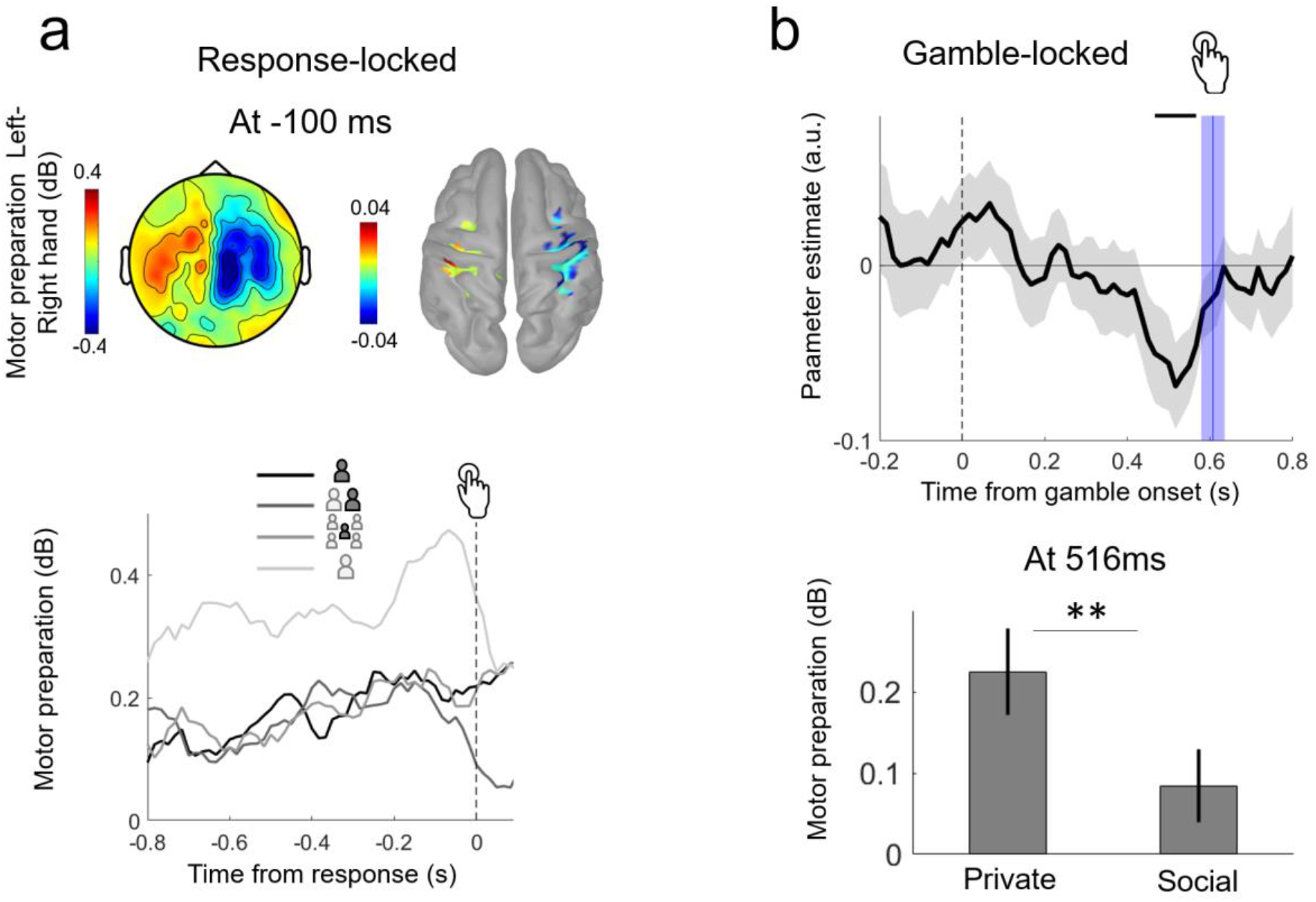
Motor preparation signals are modulated by the responsibility context. a) Topography showing the mean power of 8-32 Hz frequencies at 100 ms before the motor response for conditions where participants answered with the left hand minus the right hand and the associated estimated sources. Bottom panel: Response-locked motor lateralization: motor preparation from 0.8 seconds before and up to response measured with the power of 8-32 Hz frequencies in ipsilateral minus contralateral electrodes relative to the hand pressed, locked to the motor response in all four different contexts. b) Stimulus-locked motor lateralization. Top panel: Parameter estimate of the regression of motor preparation signal against the Social vs Private regressor. Negative parameter estimates indicate lower motor preparation in the Social vs Private context, which is significant around 500 ms after gamble onset. The black line indicates time points where the parameter estimate is significant against zero (cluster, one-tailed, pcorr<0.05). The blue bar indicates the time when the button was pressed based on reaction times and their standard errors. Bottom panel: Motor preparation signal at the peak of the effect at 516 ms for the Private vs the Social context. ** p<0.01

To test **Hypothesis 3**, we first asked whether motor preparation signals locked to gamble onset varied parametrically with responsibility. This first regression revealed a weak effect, peaking at 516 ms (T_39_=-1.90, p=0.03 one-tailed), which did not survive cluster multiple comparison corrections (p>0.22). As our hypothesis involves social contexts where responsibility was shared, we conducted a new regression that pooled the Dyadic and Group contexts (i.e. Social contexts), allowing a comparison of the Private and Social contexts. The parameter estimate of this regression was significant at ~500 ms after gamble stimulus onset (peak of parameter estimate at 516 ms, T_39_=-2.81, p=0.007, μ =-0.068, ci=[−0.11, −0.02], d=0.44, cluster from 466 ms to 566 ms, one-tailed pcorr<0.05; **Figure 4b,** top panel), where a stronger motor preparation signal was evident for Private compared to Social contexts (Direct t-test at the peak 516ms between Private and Social contexts, T_39_=2.78, p=0.008, μ=0.14, ci=[0.03,0.24], dz=0.45; **Figure 4b,** bottom panel). No significant cluster was observed for the same analyses locked to motor response rather than gamble onset (p>0.1), suggesting that the effect is related to the motor intention locked to the stimulus, rather than the motor action itself.

#### Neural correlates of increased claims of credit in Social contexts

Earlier, we provided behavioural analyses that showed that a Social (vs Private) context boosted the credit claimed for positive outcomes. In an exploratory analysis, we examined the neural correlates of this specific positive boost, focusing on Private and Social contexts for trials where the collective choice matched a participant’s vote. Concentrating on a post-outcome, 200-300 ms time window, we identified MEG signals locked to *positive* outcomes and then ran a GLM with Social vs Private as the regressor. This revealed a significant cluster (cluster alpha <0.05) that included frontal, temporal and central electrodes (see full list of electrodes in *Supplementary materials*, **Figure 5a**). No significant clusters appeared for the same regression analysis for MEG signals locked to *negative* outcomes. The same regression conducted at MEG source level for signals locked to *positive* outcomes revealed source estimates in the orbitofrontal cortex and temporal lobe, including the superior temporal sulcus (see details of brain regions in Table 2; **Figure 5b**). In the pre-central cortex, a negative parameter estimate was observed, reflecting higher activity in the Private context compared to the Social context, which is consistent with the effects of responsibility observed in this region.

**Figure 5.**
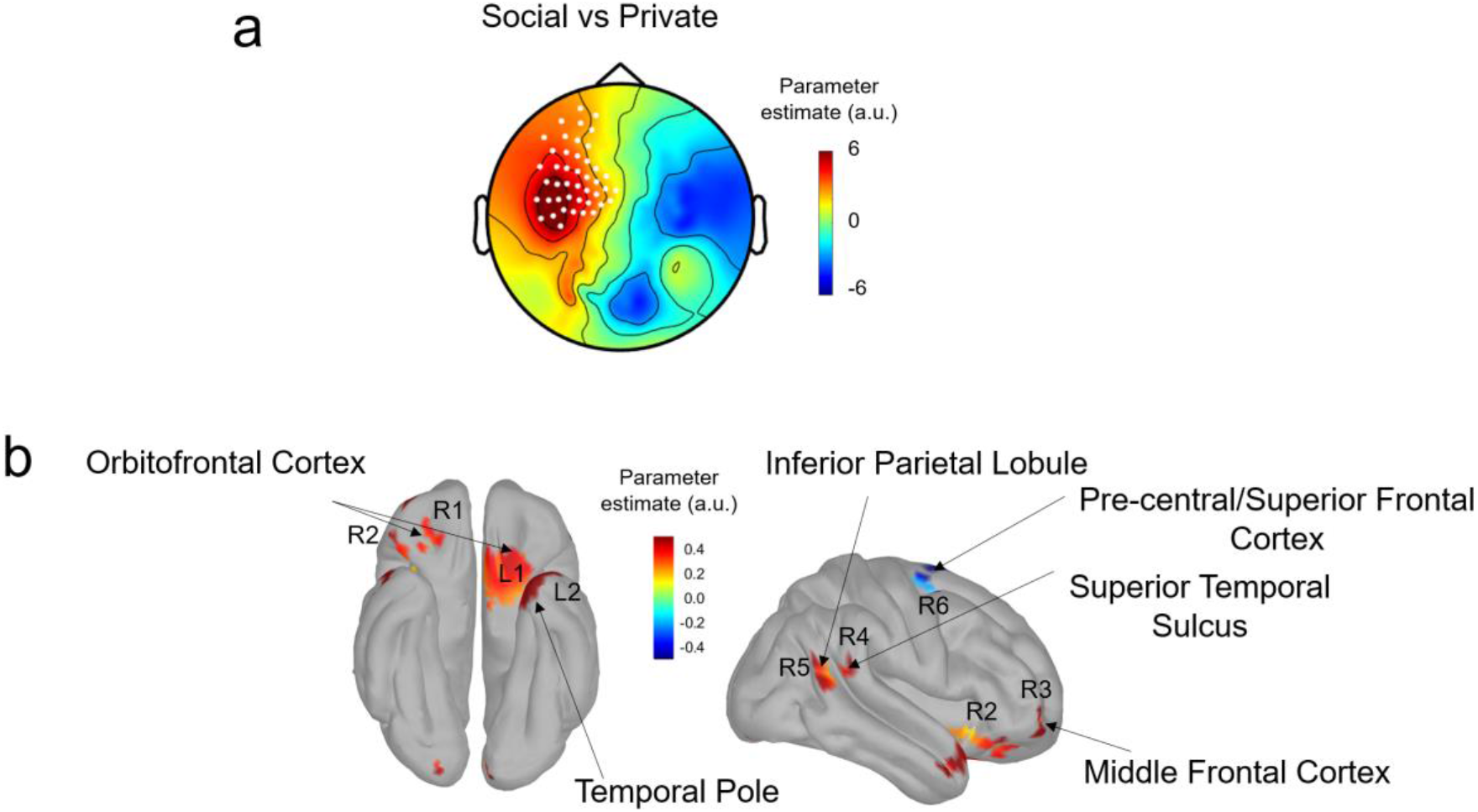
The processing of positive outcomes in Social vs Private contexts. a) Scalp topography of the parameter estimate of the regression Social vs Private on the mean MEG activity at 200-300 ms after positive outcomes following decisions that matched the participant’s vote. White dots represent the significant cluster of electrodes differentiating between Social and Private outcome processing using cluster corrections of the effect against zero (cluster alpha <0.05). b) Estimated cortical sources of the Social vs Private parameter estimate at 200-300 ms. Parameter estimates that are significant at p<0.01 are shown. R=Right hemisphere and L=Left Hemisphere, and the associated numbers refer to the different brain regions reported and anatomically localised in Table 2.

**Table 2.** Anatomical sources of Social vs Private processing of positive outcomes. Regions were determined based on the Destrieux Atlas and implemented in Brainstorm. Then, MNI coordinates were extracted from Brainstorm and projected onto the BNA atlas. Grey areas represent MNI coordinates that could not be matched to the BNA atlas.

|  | Destrieux Atlas |  |  | MNI coordinates |  |  | Human Brainnetome Atlas (BNA) |  |  |  |
| --- | --- | --- | --- | --- | --- | --- | --- | --- | --- | --- |
| Regions | Index | Name | Localisation | X | Y | Z | Label ID | Gyrus | Anatomy | Lobe |
| Left 1 (L1) | 24 | Orbital Gyri | Ventral aspect of the frontal Lobe | -21 | 14 | -27 |  |  |  |  |
|  | 63 | Medial orbital sulcus | Ventral aspect of the frontal Lobe | -14 | 17 | -15 | 49 | Orbital Gyrus | Area 13 | Frontal |
|  |  | Gyrus Rectus | Medial aspect of the frontal Lobe | -3 | 11 | -23 |  |  |  |  |
|  |  | Subcallosal Gyrus | Limbic Gyrus | -4 | 6 | 16 |  |  |  |  |
| Left 2 (L2) |  | Temporal pole | Superior aspect of the temporal lobe | -22 | 13 | -43 |  |  |  |  |
| Left 3 (L3) |  | Triangular part of the inferior frontal gyrus | Main frontal gyri | -51 | 45 | 5 |  |  |  |  |
|  |  | Inferior Frontal sulcus |  | -40 | 44 | 2.0 | 21 | Middle Frontal Gyrus | Ventral area 9/46 | Frontal |
|  | 15 | Middle Frontal Gyrus |  | -49 | 49 | 3 |  |  |  |  |
| Left 4 (L4) | 28 | Postcentral Gyrus | Lateral aspect of the parietal lobe | -24 | -31 | 77 | 161 | Postcentral Gyrus | Area 1/2/3 | Parietal |
| Left 5 (L5) | 27 | Superior parietal lobule | Superior parietal lobule | -37 | -52 | 66 |  |  |  |  |
| Right 1 (R1) | 64 | Orbital Sulcus | Ventral aspect of the frontal Lobe | 23 | 39 | -22 | 46 | Orbital Gyrus | Lateral area 11 | Frontal |
| Right 2 (R2) |  | Orbital Gyri |  | 43 | 25 | -18 | 44 | Orbital Gyrus | Orbital area 12/47 |  |
| Right 3 (R3) |  | Orbital Gyri |  | 43 | 57 | -11 |  |  |  |  |
|  | 15 | Middle Frontal Gyrus | Main frontal gyri | 39 | 63 | -1 | 28 | Middle Frontal Gyrus | Lateral area 10 | Frontal |
| Right 4 (R4) | 72 | Superior temporal sulcus | Lateral aspect of temporal lobe | 67 | -46 | 15 | 144 | Inferior parietal Lobule | Rostroventral area 39 | Parietal |
| Right 5<br>(R5) | 38 | Middle temporal Gyrus |  | 64 | -54 | 15 | 144 |  |  |  |
|  | 25 | Angular gyrus | Inferior parietal lobule | 63 | -56 | 19 | 144 |  |  |  |
| Right 6 | 69 | Superior part of the precentral sulcus | Main frontal sulci | 30 | -4.5 | 64 | 8 | Superior frontal gyrus | Dorsolateral area | Frontal |

## Discussion

We examined the behavioural and neuronal signatures of shared responsibility in collective decision-making. Behaviourally, we showed that responsibility was reduced in collective contexts compared to private, individual decision-making, and this decrease was dependent on the size of the collective. Previous neurobiological findings on responsibility for socially executed actions have consistently shown decreased outcome processing under coercion [6], cooperative gambling [14], and in the presence of another person [13]. Our work goes beyond those studies in several important aspects. First, our study examined collective *choice* rather than action. Second, we developed a systematic, parametric design with four levels of responsibility that produced a highly reliable empirical framework for studying the subtle concept of responsibility. Third, we studied the neuronal mechanisms underlying the flexible interaction between social context and outcome valence that permitted participants to cherry-pick the level of responsibility that best served them with regards to claiming more credit for positive outcomes. Finally, whereas previous neural studies on responsibility had focused exclusively on outcome evaluation [6,9,13–15], we could examine the neural substrates of responsibility during deliberation and action preparation *before* a choice was made or outcomes were known.

Our findings showed that people’s subjective reports of responsibility varied according to social context, with greater responsibility reported when making decisions privately compared to when making decisions with others. This is in line with previous studies that show that people feel less control, responsibility and regret when acting with others [3,14,16,17]. Reported responsibility varied with group size, with more responsibility reported in Dyads vs in a Group of five. This finding builds on previous results showing that people take less credit when they are problem-solving in a larger group [32].

We showed that at 200-300 ms after an outcome, MEG-recorded activity of bilateral fronto-parietal brain regions decreased linearly, from its highest at full responsibility in the Private context, to shared responsibility in Dyads, to shared responsibility in Groups, to no responsibility in the Forced context. This complements previous findings of decreased outcome processing associated with low responsibility contexts [6,13,14], but goes beyond those earlier works by showing a parametric—and not categorical—relationship to an incremental manipulation of responsibility. Second, using source estimation, we confirmed our key neural hypothesis that a marker of responsibility is localised to brain regions previously associated with a sense of agency in pre-central and post-central cortices, and superior and inferior parietal lobules [7,20–22,33]. Our findings are also consistent with previous work that identified correlates of motor intentions in the parietal cortex [7] and motor control in inferior parietal lobule [21].

It is important to note that the outcome-dependent neural signature of agency identified here (Fig. 3) was not influenced by outcome valence. In our design, we were mindful to ensure that the probability of winning did not depend on the gambling choice. The probability of winning was also independent of whether a participant decided privately, did not decide at all, or when the participant’s choice matched that of the group or not. Furthermore, to minimize incidental learning, we used a large set of visual stimuli (i.e., 40 illustrations of various gambling devices) and had the participants choose between randomly sampled pairs.

With regards to collective decision-making, several non-monetary motivational factors also came into play in our paradigm. The first factor was autonomy and control. Rewards have a higher salience when we are instrumental in obtaining them compared to when they are merely thrust upon us. Our key neural findings (**Figure 3**) are consistent with this, showing that the participant’s level of involvement in an outcome modulates an outcome-dependent neural signal. Thus, brain responses to outcomes were stronger when subjects decided privately and parametrically decreased as responsibility decreased. The second factor was the approval of others. Previous work shows that others ’approval is, inherently, capable of driving the brain’s motivational reward network even when no monetary reward is at stake [34]. In our paradigm, participants ’choices were sometimes agreed with, and other times overruled by the collective. Accordingly, we found that approval by the collective was associated with increased neural activity in the orbitofrontal cortex (OFC), the superior temporal sulcus (STS) and the temporal pole, brain regions variously associated with reward, social processing and mentalizing [35,36]. In addition to being associated with monetary reward and value processing [37,38], the human OFC is implicated in individual differences in conformity and reaction to other people’s opinions [39]. As outcome processing of a social decision involves considering other people’s responsibility, the involvement of the STS is in line with the finding that this region is involved when participants consider others ’responsibility [25].

Previous studies of the neural basis of responsibility have invariably focused on the outcomes of decisions and actions. Our study breaks with this tradition and examines neural substrates of responsibility prior to an outcome, when deliberation and action preparation are taking place. We show that the motor preparation (to pick the right or left gamble) around 500 ms after the onset of a visual display of a gambling option was reduced under shared (Social) as compared to full responsibility (Private) conditions. These results were observed only when the analysis was stimulus-locked, but not locked to the motor response, indicating that the decreased motor preparation in Social vs Private contexts most likely reflects a deliberative rather than motor process. This is also in line with findings that self-responsibility, as compared to shared responsibility, recruits brain areas associated with action simulation, including the premotor context, suggesting that higher-order social processes may relate to simple goal-directed action [25]. This effect did not follow a parametric pattern, as it did not decrease based on group size and thus seems to be related to a more general social context where responsibility is shared with others. A recent theoretical model proposed that a decreased sense of agency in social contexts relates to mentalizing processes, as people need to take into account the perspectives of others [13,40]. This model posits that through mentalizing, social contexts increase decision disfluency and action planning. Here we provide the neural evidence that action planning is indeed reduced when people make decisions with others in social contexts.

The higher responsibility ratings for positive vs negative outcomes (Fig. 2b) replicates the “self-serving bias” effect, whereby participants take more credit for positive (vs negative) outcomes [27,30,31]. It is also in line with an increased sense of agency for positive vs negative outcomes [31,41] (although the opposite effect was found in an unpredictable environment [42]). Our experimental design included a subset of trials where we probed responsibility *before an* outcome was declared (Fig 2c-d). This allowed us to treat these trials as a baseline and evaluate if the observed self-serving bias came from claiming more credit for positive outcomes and/or offloading blame for the negative ones. In addition, our design also distinguished between trials in which the group decision was in line with that of the participant and those when they were not. Consequently, collective decisions in which the group and participant agreed offered a particularly informative situation, as an individual retained *some* control but still shared responsibility with others. In these situations, participants were particularly inclined to claim disproportionately more credit for positive outcomes. These findings point to a potential motivation to join groups, particularly as claiming credit for success has been shown to increase self-esteem [30]. Interestingly, in these trials, we did not observe any offloading of blame onto others for negative outcomes. This observation is consistent with a similar recent report [40].

Our novel experimental design manipulates levels of responsibility for the outcomes of decisions and shows that responsibility influences how these outcomes are processed at 200 ms in brain regions that are related to a sense of agency. Our results are also the first to provide an outcome-independent neural signature of responsibility evident in the reduction in pre-outcome motor preparation signatures at 500 ms in shared responsibility contexts, i.e., social contexts. The finding that prospective and retrospective responsibility in social contexts involves neural mechanisms common to a sense of agency can potentially advance our understanding of the complex notion of societal responsibility and is relevant to a wide range of societal domains, including the legal and medical sectors as well as ethical issues related to artificial intelligence.

## Methods

### Participants

Forty-six healthy adults (24 females, mean age=24.13±4.42) participated in a magnetoencephalography (MEG) study, with two participants excluded due to technical errors in saving the data or in the triggers ’information. Four participants were excluded due to noisy MEG data noted in the pre-processing phase (their data had more than 10 noisy channels and/or more than 15% of trials removed after visual inspection). This left a total of 40 participants (21 females, age=24.00±4.46). The study was conducted at the Wellcome Centre for Neuroimaging, 12 Queen Square, London, WC1B 5JS. Participants were recruited by email via the University College London (UCL) SONA platform and the Institute of Cognitive Neuroscience (ICN) participants ’pool. All participants were aged 18-36 years, right-handed, with normal or corrected-to-normal vision, and had no neurological or psychiatric history. They provided written, informed consent according to the regulations approved by the UCL research ethics committee (Project ID Number 9923/002). Participants were informed that they will receive £25 for their participation and a bonus of up to £5 based on their gains during the experiment. All participants were given the bonus and compensated with £30.

### Experimental design

Stimuli were generated using Cogent 2000 and Cogent Graphics toolboxes running in MATLAB (MathWorks, Na-tick, MA, USA). The task was presented as a learning game where participants had to choose between two different gambles that supposedly had different probabilities of getting a positive or negative outcome. The two gambles corresponded to two different images of hand-drawn, realistic gambling devices from among 40 total drawings (an example of two drawings is shown in **Figure 1b**). Two gambles were pseudo-randomly presented for each trial from among a set of 10 gambles for each block: We controlled that the gambles were presented for approximately the same amount of repetition within each context (contexts described below **Figure 1a**) and each block, and not more than three times in a row. Unbeknownst to participants, the gambles were not associated with different probabilities that yielded positive or negative outcomes, as the frequency of positive and negative outcomes was entirely controlled for.

In each trial, following a fixation cross displayed for 700-900 ms, participants first saw a cue (duration 1000 ms) indicating which context they would be playing in. There were four possible contexts (**Figure 1a**): (1) **Private**; (2) **Dyadic:** A participant plays with one other player, so that both players make a decision, but only one of their decisions is selected to determine the outcome; (3) **Group:** A participant plays with four other people where the selected gamble is based on a majority vote (the gamble picked by three or more people)—(2) and (3) are referred to as **Social** contexts; and (4) **Forced**: Another player chooses on behalf of the participant (participants did not have the choice to *not* accept the other player’s selection). After the cue, a fixation cross was displayed for a variable period of 1000-1200 ms. Then, two gambles appeared on the screen—one on the left and one on the right side of the cross—and participants had to select between them by pressing the respective button on the devices that they held in their right (Right gamble) and left (Left gamble) hands (**Figure 1b**).

In the Forced context, participants were instructed to always press the same button after seeing the gambles. They were informed whether to press the left or the right button (constant throughout the experiment) in this context at the beginning of the experiment, and this was counterbalanced across participants. This button press was included to ensure that action requirements for all trial types were identical, but does not indicate an actual choice, which allowed us to maintain identical stimulus-response mappings. Participants were told that different gambles had different probabilities of winning and losing, and that they should try and pick the one that had a higher chance of winning. In all trials, the response window was two seconds, otherwise the trial was classified as missed (even in the Forced context). The selected gamble was then displayed for 200 ms followed by a variable blank period of 650-850 ms. The trial ended with an outcome (positive/negative) presentation (1000 ms). Participants were told that one trial would subsequently be picked from each block and that they would earn a cumulative bonus based on whether the outcomes of selected trials were positive or negative, with a missed trial counting as a negative outcome. Note that positive outcomes allowed them to win a bonus, while negative outcomes were similar to ‘neutral ’ outcomes as they could not ‘lose ’any money.

In one-third of the trials, immediately after having made a decision, but *before* an outcome was shown (i.e., T1, see **Figure 1b**), participants rated how responsible they felt for the upcoming outcome (from ‘not at all ’to ‘partially ’to ‘very much ’on a continuous scale). In another third of the trials, they made a similar rating immediately *after* the outcome (i.e., T2). In one-third of the trials, no ratings were required.

Participants completed 384 trials in eight blocks of 48 trials each. Each block was composed of a balanced number of trials: 2 contexts (Alone, Forced) X 2 outcomes (Positive/Negative) X 3 scales (T1, T2 or none) X 2 repetitions; and in the Social contexts: 2 contexts (Dyadic, Group) X 2 outcomes (Positive/Negative) X 3 scales (T1, T2 or none) X 2 feedback (whether the participant’s vote was selected or not).

### Behavioural analyses

Responsibility ratings, based on a continuous scale, were z-scored for each participant before subject and group-level statistical analyses were made and are reported in arbitrary units. A general linear model (GLM) was performed for each participant’s responsibility ratings with the parametric responsibility contexts as a regressor [1,2,3,4]: 1= Private, 2= Dyadic, 3= Group, 4= Forced. The betas of these regressions were tested against zero with a t-test for the group level statistics. Paired student t-tests were used to compare ratings in each context from one another, and to assess the differences between ratings for positive and negative outcomes as well as differences between ratings both before and after an outcome. Reaction times for the gamble decision were also compared between contexts using paired student t-tests. Confidence intervals of the effects and effect size (Cohen’s d) are provided.

### MEG acquisition and pre-processing

MEG data was recorded using a 275-channel CTF Omega system whole-head gradiometer (VSM MedTech, Coquitlam, BC, Canada). Neuromagnetic signal was continuously recorded at a 600 Hz sampling rate. After participants were comfortably seated in the MEG, head localiser coils were attached to the nasion and 1 cm anterior to the left and right tragus to monitor their head movement during recording. Due to technical issues, three gradiometers were disabled from the system: MLO42, MRC12, MLT54, leaving a total of 272 instead of 275 channels. Three additional channels recorded eye movements (x, y movement and pupil diameter) using an eye-tracker (SR Research EyeLink non-ferrous infrared eye tracking system).

Stimuli were projected at a 60 Hz frequency on a screen of 42 x 32 cm, with a distance of about 60 cm between the screen and the eyes. During piloting, a photodiode placed on the screen allowed us to measure a delay of about 33 ms between the trigger signal and the projection of stimuli. The appearance of stimuli on the screen was therefore monitored with a photodiode attached at the lower edge of the screen. A black square was presented there for 50 ms at the time when the stimulus was presented. The signal from the photodiodes was recorded in parallel to the other MEG channels to provide a precise temporal marker for the appearance of stimuli. All triggers were matched to the signal of the photodiode associated with each stimulus of interest.

We used FieldTrip [43] to process the data. The raw MEG data was notched for the 50, 100 and 150 Hz power line noise before visual inspection (combined with an automatic detection of artifacts) to reject trials with jumps and strong muscular activity. The data was low-pass filtered at 35 Hz and epoched using the photodiode signal locked to gamble onset, response onset and outcome onset. Independent component analysis (ICA) was performed on the data to reject eye blinks, saccades and heartbeat components thanks to the visual inspection of the components. The mean proportion of rejected, artifacted trials was 7.7% ±3.1 for context-locked data, 6.5%±3.4 for gamble locked data, 6.4% ±3.4 for response-locked data and 5.9%±2.8 for outcome-locked data.

Time-frequency decompositions were performed by computing the spectral power of the 8-32 Hz frequency bands using multi-tapering transform (Slepian tapers, 8-32Hz, four cycles) centred on gamble presentation and response onset, using the FieldTrip function ft_freqanalysis. The power spectrum was extracted for the main analysis, and the complex Fourier spectrum was extracted for the source reconstruction analyses.

### MEG analyses

#### Regression analyses of outcome-locked MEG signals

We performed single-trial regressions of MEG signals against variables of interest:

- Responsibility context [1,2,3,4] as for the behavioural analyses: 1= Private, 2= Dyadic, 3= Group, 4= Forced. Please note that negative parameter estimates indicate higher activity for more responsibility.
- Social (+1 for the Dyadic and Group contexts) vs Private (−1) contexts.

As our hypothesis predicted that outcome processing would be modulated by responsibility at 200-300 ms, we performed regressions on the mean activity of MEG signals at 200-300 ms after outcome onset at each of the 272 electrodes. Beta coefficients of regressions for each participant were tested against zero for the group level statistics. Multiple comparison corrections were performed in the electrode space using ft_prepare_neighbours in FieldTrip, coupled with non-parametric Montecarlo statistics to determine the clusters of electrodes where these regressions were significant with a p-value<0.05 (cluster corrections, cluster alpha=0.05, test statistic set as the maximum level of the cluster-level statistic, alpha=0.05, 10000 randomizations, 2 minimal neighbouring channels).

#### Motor preparation measures in the time-frequency domain

##### Motor lateralization

The suppression of 8-32 Hz frequency bands in the hemisphere contralateral to the hand used in a motor press response provides a neural marker for motor preparation [26]. For each participant, the power of this frequency band when participants responded with their right hand was subtracted from the power when they responded with their left hand, at 100 ms before a response press. When averaged across participants, this allowed us to identify the central electrodes with maximal suppression (**Figure 4a**). Motor lateralization for responses with the right and left hand was obtained by subtracting power activity in the central electrodes contralateral to the utilised hand from power activity in the central electrodes ipsilateral to the hand used, thus resulting in positive motor preparation as shown in **Figure 4b** (top panel). Right electrodes: ‘MRC13’; ‘MRC14’; ‘MRC22’; ‘MRC31’; ‘MRC32’; ‘MRC41’; ‘MRC42’; ‘MRC53’; ‘MRC54’; ‘MRC55’.

Left electrodes: ‘MLC13’; ‘MLC14’; ‘MLC22’; ‘MLC31’; ‘MLC22’; ‘MLC41’; ‘MLC42’; ‘MLC53’; ‘MLC54’; ‘MLC55’.

We performed regressions on this motor lateralization measure at each time point, locked to the gamble and to the response, using the following regressor:

- Responsibility context [1,2,3] 1= Private, 2= Dyadic, 3= Group
- Social context (+1 pooling both Dyadic and Group) vs Private (−1)

We added reaction times (z-scored) as an additional regressor, to account for effects over and above differences in reaction times for motor press. The parameter estimates were tested for significance against zero at each time point from 0 to 600 ms after gamble onset, with multiple comparison corrections across time points implemented using non-parametric cluster-level statistics [44]. The pairing between experimental conditions and MEG signals was shuffled 1,000 times, and the maximum cluster-level statistics (the sum of one-tailed t-values across contiguously significant time points at a threshold level of 0.05) was extracted for each shuffle to compute a null distribution of the size of the effect across a time window of [0,+600] ms locked to stimulus presentation, or [-600,0] ms locked to response onset. For significant clusters in the original (non-shuffled) data, we computed the proportion of clusters in the null distribution where statistics exceeded that of the one obtained for the cluster in question, as it corresponds to its ‘cluster-corrected ’p-value.

#### Source reconstruction

Minimum-norm source estimates were performed using Brainstorm [45]. We computed Kernel inversion matrices for each subject and for each of the eight blocks (recommended due to differences in participants’ head movements in different blocks), using all trials consisting of non-overlapping time windows locked to context, gamble and outcome. We used a generic brain model taken from the default anatomy in Brainstorm: ICBM 152 Nonlinear atlas version 2009 [[46,47]. The head model was computed with an overlapping spheres method. The noise co-variance was computed based on a 900 ms baseline before the context onset (i.e. the baseline of the whole trial). Sources were computed with minimum norm imaging and the current density map method. Following recommendations for when there is no individual anatomical IRM available, we chose unconstrained solutions to source estimation. In unconstrained source maps, there are three dipoles with orthogonal orientations at each cortex location (15002 vertices X 3 orientations = 45006 × 272 electrodes, 8 Inversion matrices—for each of the 8 blocks—for each participant). To display these as an activity map and perform statistics, the norm of the vectorial sum of the three orientations at each vertex is computed as follows:

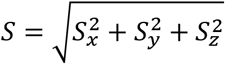

##### GLM in the source space

At 200-300 ms after outcome onset for each subject, the mean MEG signals were multiplied by individual Kernel matrices for each block, and the norm was computed before implementing the GLMs at each vertex at the source level. Finally, the betas at each vertex were averaged across blocks, resulting in one map of parameter estimates for the effect of interest for each participant. Then, for group level analyses, t-tests against zero were done at each vertex, and only the vertices with mean parameter estimates significant across participants at p-value<0.01 are shown. To assess the anatomical location of the significant vertices, we first reported regions based on the Destrieux Atlas [48] that is provided in Brainstorm. Second, we extracted MNI coordinates of regions and projected them onto the human Brainnetome Atlas (BNA) [49] (through MRICron where MNI coordinates can be matched to the anatomical location of the chosen atlas, NITRC: MRIcron: Tool/Resource Info).

##### Motor lateralization

At 100 ms before response press for each participant, the MEG Fourier transforms from the time-frequency decomposition analysis were multiplied by individual Kernel matrices for each frequency and trial. Then, after taking the power of the magnitude of the complex Fourier spectrum, a mean was performed on the frequencies (8-32Hz) and the conditions where the participant responded with the right hand and the conditions where the participant responded with the left hand. The sum of the power for each orientation was computed for each block, before doing a mean on the eight blocks to obtain one activity map per condition and per subject. Left press vs Right press were then contrasted by performing a t-test in the source level after a z-score of the activity map per participant, and keeping only significant vertices at p<0.01 for the figure. The mean difference across all participants at the vertices with this significant effect is shown (**Figure 4a**, bottom panel).

## Acknowledgments

This project has received funding from the European Research Council (ERC) under the European Union’s Horizon 2020 research and innovation programme (PI: BB grant agreement No. 819040 - acronym: rid-O) and from the Wellcome Trust (PI: RJD grant number 098362/Z/12/Z). MEZ was funded by the Wellcome Trust, grant number 204702, and is currently supported by the European Union’s Horizon 2020 Marie Curie Individual fellowship, grant number 882936 acronym SIND. BB was supported by the Humboldt Foundation and by the NOMIS foundation. RJD is in receipt of a Wellcome Trust Investigator Award.

We would like to thank Daniel Bates for his valuable help with data acquisition, the MEG team at the Wellcome Trust Center for Neuroimaging for their feedback on the project, David Wurzer for his help with data pre-processing, Alizee Lopez for her help with source analyses, and Stephanie Don for editing the manuscript.

## Author contributions

M.E.Z and B.B designed the study. M.E.Z programmed the experiment, and collected and analysed the data. M.E.Z, R.D and B.B discussed analyses and interpreted the results. M.E.Z and B.B wrote the paper and R.D provided important revisions.

## Materials & correspondence

Correspondence and material requests should be addressed to Marwa El Zein.

## Data and code availability

MATLAB code for the analyses of the behavioural and neural data and the behavioral data will be shared on the open science framework upon publication. Further requests can be addressed to M.E.Z.

## Supplementary materials

### Behavioural results

Change in reported responsibility from before to after outcome in all conditions.

**Supplementary Figure 1.**
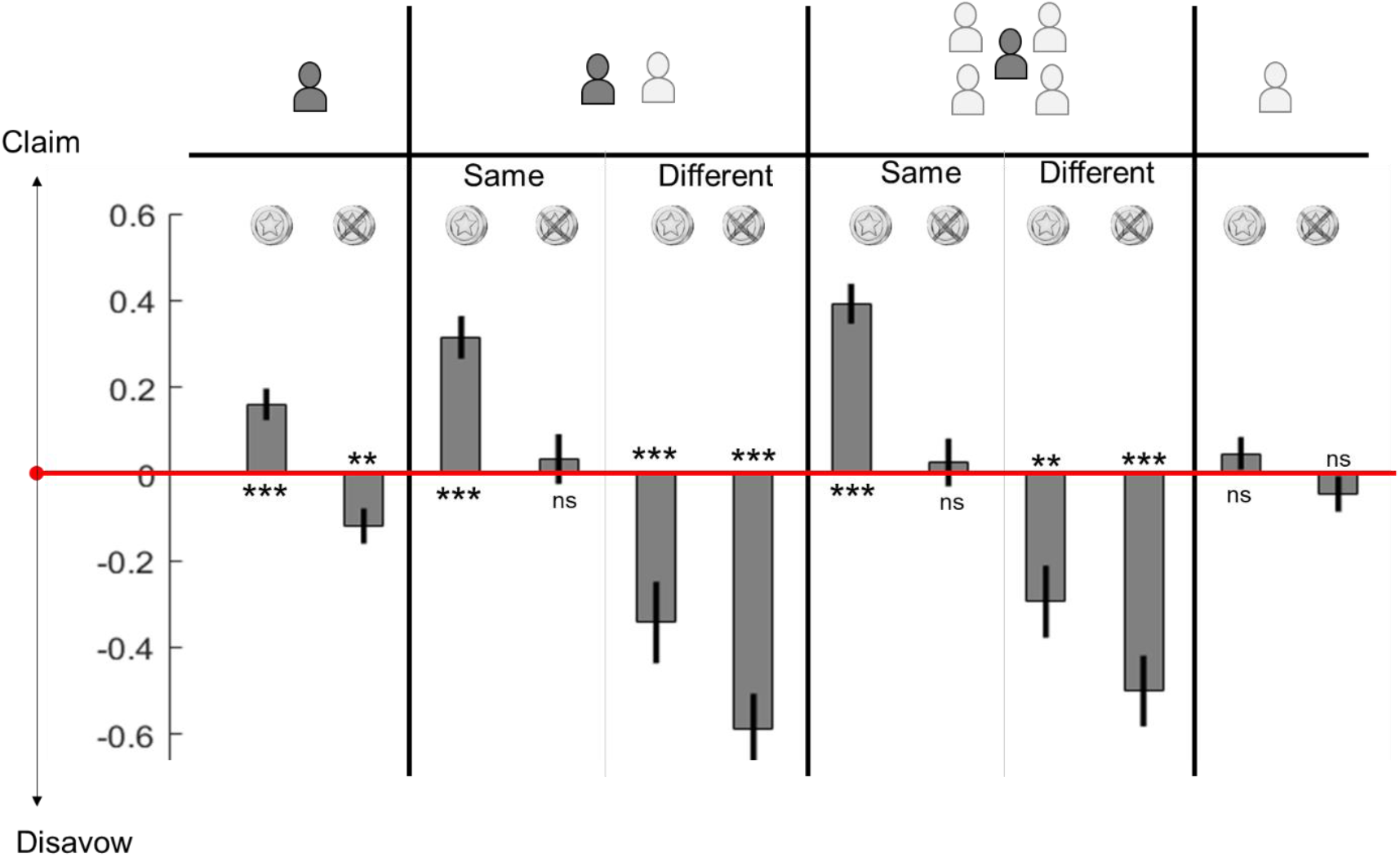
Y-axis shows the difference between responsibility ratings after vs before an outcome. Error bars represent standard errors. ***: p<0.001, **: p<0.01, ns: non significant.

### MEG results

#### • Regression of MEG signals, outcome-locked, at 200-300 ms against responsibility levels

Two clusters of electrodes showed significant activity which survived correction for multiple comparisons (cluster alpha < 0.05, **Figure 3a**, right panel): These electrodes match the expected fronto-parietal network as they consist of **central** (MZC04, MZF03, Left MLC11 to MLC17, MLC21 to MLC25, MLC31, MLC32, MLC41; Right MRC11 to MRC17, MRC22 to MRC25, MRC31, MRC32 MRC41 MRC42, MRC52 to MRC55, MRC61 to MRC63), **frontal** (Left MLF35, MLF44 to MLF46, MLF53 to MLF56, MLF61 to MLF67; Right MRF46, MRF55, MRF56 MRF64 to MRF67) and **parietal** electrodes (Left MLP45 and MLP57, Right MRP11 MRP12 MRP21 to MRP23, MRP32 to MRP35, MRP43 to MRP45, MRP56, MRP57), as well as four **temporal** electrodes (MLT12, MLT13, MRT13, MRT14).

#### • Regression of MEG signals, outcome-locked at 200-300 ms, against Social vs Private

A significant cluster (cluster alpha <0.05) was observed that included **frontal** (MLF12 to MLF14, MLF22 to MLF25, MLF32 to MLF35, MLF43 to MLF46, MLF53 to MLF56, MLG63 to MLF67), **temporal** (MLT12 MLT13, MLT21 to MLT23), and **central** (MLC12 to MLC16, MLC21 to MLC24, MLC31 MLC41 MLC51 MLC52) electrodes (**Figure 5a**).

#### • Lateralized motor preparation results

Regression of the motor preparation signals at each time point locked to the response, with the Private+Social vs Forced as a regressor, showed that the Forced context significantly differed from Private+Social at ~50ms before response (T_39_=2.86, p=0.006, μ =0.10, ci=[0.03,0.17], dz=0.43, cluster from 100ms to response, one-tailed pcorr<0.05). This is coherent with the fast and known motor response in the Forced context. This effect was not observed when doing the same analyses stimulus-locked.

**Supplementary Figure 2.**
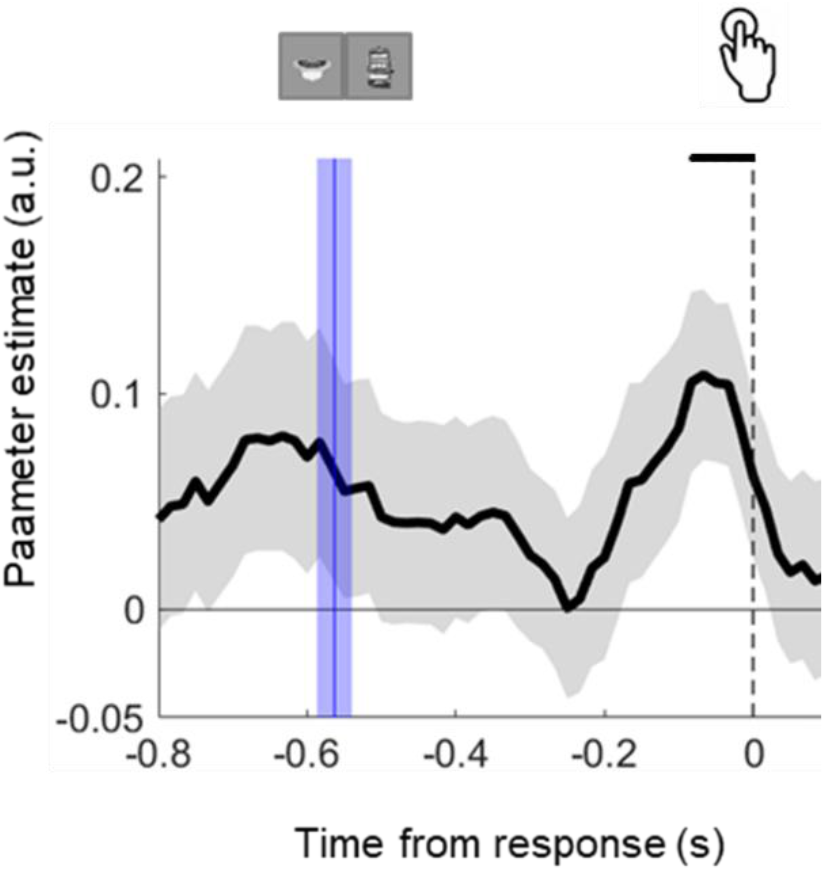
Response-locked motor lateralization. Parameter estimate of trial-by-trial regression of motor preparation against [Forced +1] vs [Private + Social contexts −1] regression in the motor lateralization measures. Positive parameter estimates indicate higher motor preparation in the Forced context: Motor preparation is higher when the response is known in the Forced context. Dots indicate time points where the parameter estimate is significant against zero (cluster one-tailed pcorr p<0.05). The blue bar indicates the time when the stimulus appeared based on mean reaction times and their standards of error.

